# A gene regulatory network for apical organ neurogenesis and its spatial control in sea star embryos

**DOI:** 10.1101/036624

**Authors:** Alys M. Cheatle Jarvela, Kristen A. Yankura, Veronica F. Hinman

**Affiliations:** Department of Biological Sciences, Carnegie Mellon University, 4400 5th Ave, Pittsburgh, PA 15213 USA

**Keywords:** Sea star, Patiria, neurogenesis, GRN

## Abstract

How neural stem cells generate the correct number and type of differentiated neurons in appropriate places is an important question in developmental biology. Although nervous systems are diverse across phyla, many taxa have a larva that forms an anterior concentration of neurons, or apical organ. The number of neurons in these organs is highly variable. We show that neurogenesis in the sea star larvae begins with *soxc*-expressing multipotent progenitors. These give rise to restricted progenitors that express *lhx2/9*. *Soxc*- and *lhx2/9*-expressing cells are capable of undergoing both asymmetric divisions, which allow for progression towards a particular neural fate, and symmetric proliferative divisions. Nested concentric domains of gene expression along the anterior-posterior (AP) axis, which have been observed in a great diversity of metazoans, control neurogenesis in the sea star by promoting particular division modes and progression towards becoming a neuron. This work, therefore, explains how spatial patterning in the ectoderm controls progression of neurogenesis. Modification to the sizes of these AP territories provides a simple mechanism to explain the diversity of neuron number found among apical organs.

**Summary Statement:** The progression of apical organ neurogenesis in the sea star is controlled by regulatory anterior-posterior patterning domains.

## Introduction

Knowledge of how stem cells form distinct, differentiated neural cell types in the correct place, time, and numbers is important for understanding how nervous systems develop, function, and repair. The specification of neural cell fate involves a complicated combination of temporal-identity factors and positional information that direct the changing competence of the progenitor states (Kohwi and Doe, 2013). Together, these types of information instruct multipotent neural progenitors to first become restricted progenitors of a particular type, and later to execute a particular terminal differentiation program (Guillemot, 2007).

Much of the work towards understanding neurogenesis has utilized either vertebrate animals or invertebrate protostomes, namely *Drosophila melanogaster* and *Caenorhabditis elegans*, as model systems. Vertebrates, however, are exceptional among metazoa, due to their highly complex nervous systems that contain astonishingly large numbers of neurons and neuronal cell types. The genetic programs that control neurogenesis are therefore extraordinarily difficult to comprehensively study and understand in vertebrate systems. For this reason, many lines of research have been dedicated to understanding neurogenesis in invertebrate models. However, there are some fundamental differences in neurogenesis between invertebrate models and commonly used vertebrate models. For example, in *Drosophila*, neurogenesis proceeds as an invariant chain of asymmetric divisions, while the radial glia of vertebrates have more flexibility in their mode of division, allowing for more proliferative cells divisions and ultimately more neurons (Price et al., 2011). These differences make it unclear which features of the neurogenesis genetic programs are common to bilateria, and which might be tied to increase in numbers and diversity of neurons in vertebrates. Knowledge of the gene regulatory networks (GRNs) that control neurogenesis in a broader sampling of invertebrate taxa, especially invertebrate deuterostomes, which are closely related to vertebrates, is needed to understand basal mechanisms of neurogenesis and how complexity arose in vertebrates.

The larvae of echinoderms offer such a deuterostome invertebrate model. As adults, echinoderms have a radial nervous system reflecting their derived, pentaradial symmetry, which is difficult to relate to nervous systems of other deuterostome phyla (Holland et al., 2013). However, echinoderm larvae, which are bilaterally symmetrical, have neurons associated with ciliary bands that transverse the ectoderm, as well an anterior localization of serotonergic neurons, called the apical organ (Bisgrove and Burke, 1986; Chee and Byrne, 1999; Nakajima et al., 2004b). Many marine larval forms, including those of other deuterostomes, such as the tornaria larvae of hemichordates (Nakajima et al., 2004a), and also protostomes such as mollusks (Kempf et al., 1997), and annelids (Marlow et al., 2014), form apical organs. These apical organs exhibit diversity in their size and morphology. Cnidarian embryos lack an anterior concentration of serotonergic neurons, but have a molecularly homologous territory with sensory function, suggesting a common origin of metazoan apical organs and a potentially ancient gene regulatory process (Sinigaglia et al., 2013). Because apical organs are found in such diverse organisms, they offer a potentially homologous neural structure which is simple enough to dissect at the GRN level. While apical organ territory patterning is relatively well-studied and conserved (Range, 2014), the process of neurogenesis that directs an ectodermally-derived progenitor cell to progress to a differentiated serotonergic neuron in this territory is still poorly understood in any of these taxa.

Previously, we began to characterize neurogenesis in the bipinnaria larvae of the sea star (Yankura et al., 2013). We also found that the ectoderm of this organism is initially broadly neurogenic, as it expresses the extremely well-conserved neural stem cell transcription factor, *soxb1* (Miyagi et al., 2009), throughout this territory in early development (Yankura et al., 2013). *Soxc*-expressing (*soxc*+) cells are then partitioned from this ectoderm through Delta-Notch signaling (Yankura et al., 2013). Knock-down of Soxc leads to the loss of all neural cells types, and potentially also other cell types. Therefore, we predicted that these *soxc*+ cells are likely to be multipotent progenitors (Yankura et al., 2013). Intriguingly, *soxc*+ cells are found throughout the ectoderm, but differentiated neurons only form in the ciliary bands and apical organ. Therefore, we sought here to understand how anterior-posterior (AP) patterning could direct pan-ectodermal *soxc*+ cells to progress to more restricted progenitors, and finally to differentiated neurons, within a particular ectodermal territory. We focus specifically on the formation of the neurons in the APD as these most likely represent homologous structures and are clearly defined by serotonin immunoreactivity.

We first confirm that *soxc*+ cells are indeed a proliferating population of multipotent progenitors. These progenitors give rise to more restricted progenitors, which become serotonergic neurons, and are further defined by the expression of the LIM homeodomain factor, *lhx2/9*. This only occurs within the retinal homebox, *rx*, regulatory environment. These restricted progenitors differentiate into serotonergic neurons within the apical pole domain (APD) territory, which is the anterior-most region of the ectoderm. This territory is defined by the expression of the forkhead factor, *foxq2*. Therefore, surprisingly, we show that the role of AP restricted patterning genes, like *foxq2* and *rx*, is to regulate the progression of multipotent progenitors, first to restricted progenitors and finally to post-mitotic differentiated neurons, such that each step occurs in a discrete spatial domain. Patterning of the AP domains also regulates the size of the neural proliferation zone and the ultimate number of neurons in the apical organ. This presents a possible conserved basal role for these AP restricted patterns in controlling the progression of neurogenesis.

## Results

### *Soxc*+ multipotent progenitors produce *lhx2/9*+ restricted progenitors in the anterior ectoderm

The serotonergic neurons of the apical organ are found in two bilateral clusters within the anterior, dorsal ectoderm of the *Patiria miniata* larvae (Fig. 1A-B) and other sea star species at this stage (Chee and Byrne, 1999; Nakajima et al., 2004b). We previously showed that ectodermal *elav* is a marker of post-mitotic neurons, and is expressed by both the ciliary band neurons and anterior, dorsal ganglia (Yankura et al., 2013). *Elav* is also expressed within migratory mesenchyme, and thus it is likely that single sea star *elav* ortholog takes on the roles of the multiple paralogs for this gene in vertebrate animals (Yankura et al., 2013). We can however readily distinguish between the mesenchymal cells and ectodermal cells, because the mesenchyme does not embed itself within the ectoderm (SI 1F). Additionally, we had shown that correct Soxc function is needed for the ectodermal expression of *elav* in gastrula stage larvae. *Soxc*+ cells are found scattered throughout the ectoderm at 48 hours post-fertilization (h) (Yankura et al., 2013), although the differentiated neurons of the apical organ are restricted to the dorsal regions of the apical pole domain (APD). For the study here, we specifically demonstrate that some of these *soxc*+ cells comprise the cell lineage that will give rise to serotonergic neurons. We predict that other soxc+ cells will give rise to other types of neurons, including those of the ciliary band. To help demonstrate that some soxc+ cells are indeed the lineage that leads to serotonergic neurons we developed a GFP knock-in BAC for *PmSoxc*, in which *gfp* was inserted in place of the single *soxc* coding exon. When injected into embryos, the BAC becomes stably incorporated into a clonal patch of cells, which express *gfp* under the regulatory control of the *soxc* locus. As GFP protein is quite stable, it can be used to track cells that had once expressed *soxc*, even when the *soxc* transcripts are no longer being expressed. (Fig S1). Some larvae injected with this construct eventually give rise to cells with GFP label in what appear to be serotonergic neurons based on their location and morphology (Fig. 1C). We demonstrate, using immunofluorescence against GFP and serotonin, that the mosaic expression of GFP driven by this BAC construct indeed overlapped with clusters of serotonergic neurons at larval stages (Fig. 1D). This demonstrates that at least some *soxc*+ cells eventually take on a serotonergic neuron fate.

**Fig. 1.**
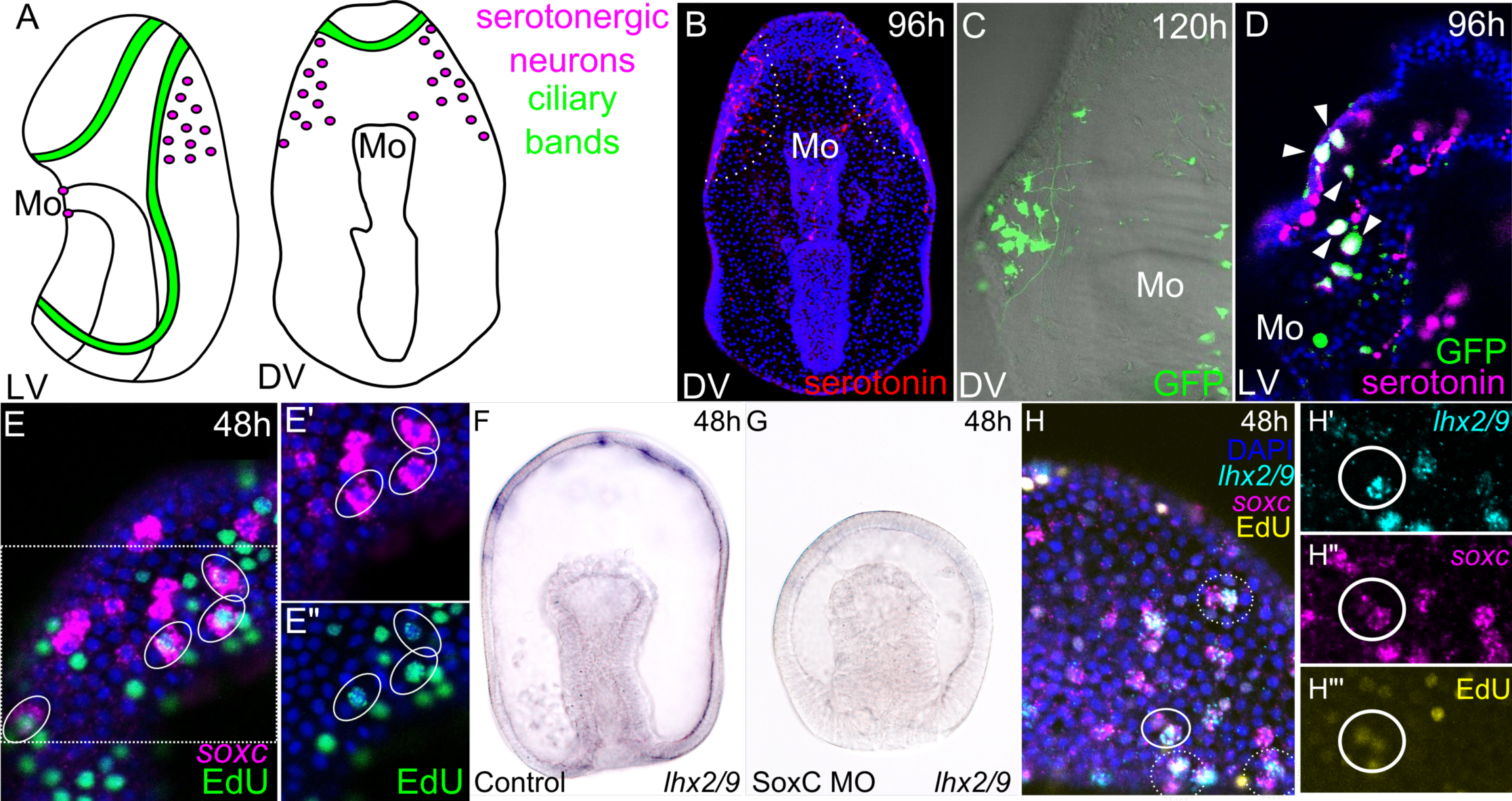
*Soxc*+cells are multipotent progenitors and generate *lhx2/9*+ restricted progenitors in the anterior ectoderm. A. Schematic of the larval nervous system, consisting of serotonergic neurons (magenta) in the mouth and apical organ as well as ciliary band neurons (green). B. Dorsal view of a 96 hour (h) sea star larvae. Apical organ neurons are stained by immunofluorescence with rabbit-anti-serotonin (red), and appear in two clusters in the anterior ectoderm. Nuclei stained with DAPI are shown in blue (B, D, E, H). Mo indicates location of mouth. C. A 120 h larvae injected with *soxc* GFP recombinant BAC construct has a labeled population of cells that have the correct position and morphology to be serotonergic neurons when compared to B. D. Lateral view of a 96h larvae injected with *soxc* GFP recombinant BAC construct, and stained by immunofluorescence for GFP and serotonin. Co-localization (indicated by arrows) demonstrates that some formerly *soxc*+ cells eventually go on to become serotonergic neurons. E-E". FISH (magenta) combined with EdU labeling (green) at 48 h results in co-labeling of *soxc*+ cells that are actively proliferating in the anterior ectoderm (circled). Region enlarged in E' and E" indicated by dashed box in E. F. *Lhx2/9* is normally expressed in spots in the anterior ectoderm of 48 h embryos. G. Knock-down of Soxc through a MO results in loss of *lhx2/9* expression. H-H"'. Double FISH plus EdU labeling reveals *soxc* expression (magenta) frequently occurs in pairs, and co-expression with *lhx2/9*(cyan) occurs in one of the two *soxc*+ cells, indicated by both solid and dashed circles. Solid circle in G indicates section enlarged in H'-H’’. H’-H’’. Asymmetric pairs are the result of recent cell division, as demonstrated by light EdU label (yellow) persisting in the pair of cells after a brief labeling period plus 30m chase (H’’’).

Given the role of *soxc* orthologs in regulating proliferation during development of neural precursors in vertebrates (Bergsland et al., 2011; Wang et al., 2013), we wanted to test whether these *soxc*+ cells are multipotent progenitors in the sea star. On closer inspection, we show that *soxc* is often expressed in a pair of adjacent cells, which appear to be recently divided cells (Fig. 1E). To determine whether these cells were dividing, we labeled proliferating cells by bathing embryos in a short pulse of EdU to identify cells in S-phase. This showed that some *soxc*+ cells, identified by fluorescent *in situ* hybridization (FISH), are indeed undergoing cell division (Fig. 1E-E”). This is consistent with a model in which *soxc*+ cells represent a population of multipotent progenitors.

We next turned our attention to *lhx2/9*, which is expressed in individual cells within the anterior-dorsal ectoderm at 48 h (Fig. S2A-B). By 96 h, *lhx2/9* is expressed in two bilateral clusters in the anterior dorsal ectoderm (Fig. S2C-D) which is reminiscent of the pattern of serotonergic neurons in 96 h larva (Fig. 1B) (Chee and Byrne, 1999; Nakajima et al., 2004b). We speculated that *lhx2/9* could be important to apical organ development and examined its expression and function more carefully. First, we show that Soxc function is needed for the formation of the *lhx2/9*+ cells in the anterior ectoderm (Fig. 1F-G), as *lhx2/9* expression is entirely absent when Soxc is knocked-down using a specific morpholino antisense oligonucleotide (MO). We examined the expression of *lhx2/9* and show that it is frequently expressed in one of the two paired *soxc*+ cells (Fig. 1H). This co-expression occurs only in the dorsal anterior ectoderm where *lhx2/9* is expressed, and hence pairs of *soxc*+ cells are completely *lhx2/9-* in the more posterior ectoderm (Fig. S2E). Because these pairs are reminiscent of recent cell division, we performed an EdU pulse-chase to label recently divided cells, which appear as pairs of adjacent cells marked by faint EdU incorporation. We are able to label some *soxc/lhx2/9*+ pairs with EdU in this way (Fig. 1H-H”’), which further confirms that *lhx2/9*+ cells are the progeny of *soxc*+ cells and that *soxc*+ cells can undergo asymmetric cell division. That is, one daughter cell maintains a stem cell state and the other, which expresses *lhx2/9*, may now be restricted to serotonergic versus ciliary band neural fate. Thus, both ciliary band and apical organ neurons originate from an ectoderm that has broad neurogenic potential, as indicated by the broad territory of *soxc* expression, but subsequent expression of *lhx2/9* may mark restriction towards apical organ neuron fate.

### *PmLhx2/9+* restricted progenitors produce the serotonergic neurons of the apical organ in the apical pole domain

We next wanted to determine whether *lhx2/9* is needed for the formation of apical organ neurons When Lhx2/9 is knocked-down, *elav* is no longer expressed in the APD of 48 h embryos (Fig. 2A-B), although we note that mesodermal elav expression appears unaffected. We had previously speculated that the *elav*+ cells in the APD at this stage will contribute to the apical organ rather than the ciliary band neurons, and that ciliary band neurons differentiate in later development (Yankura et al., 2013). In support of this, we now show that *elav* expression is specifically lost from the apical organ of 120 h Lhx2/9 morphants (Fig. 2C-D). By this stage, the neurons are normally located more posteriorly, as seen in Fig. 1B. Other domains of *elav* expression, namely the mesodermal bulb of 48 h embryos and ciliary band neurons of 120 h larvae, are unaffected by the loss of Lhx2/9, demonstrating a very specific role for this gene. As predicted by these results, Lhx2/9 morphants do not produce serotonergic neurons, although these are present in control MO injected larvae (Fig. 2E-F).

**Fig. 2.**
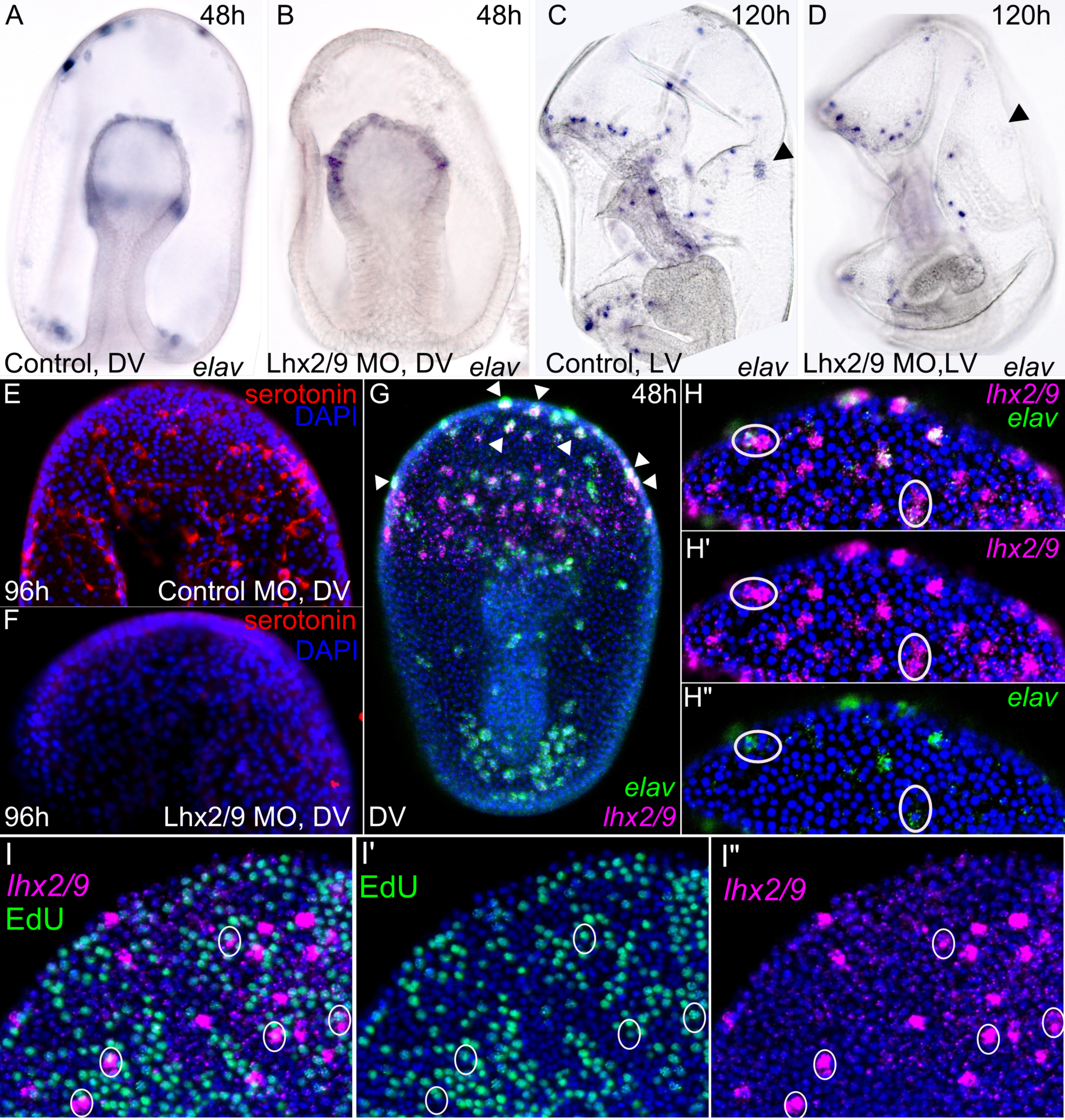
*PmLhx2/9*+ restricted progenitors give rise to the serotonergic neurons of the apical pole domain. In 48 h embryos, *elav* is normally expressed in the APD and also in the mesodermal bulb (A). Expression is lost only from the APD when Lhx2/9 is perturbed by a MO (B). Lhx2/9 is specifically needed for serotonergic neuron development. At 120 h, larvae have two neuron populations marked by *elav* expression (C). Only the apical organ *elav* expression is lost in Lhx2/9 morphant embryos (D), indicated by arrow heads. Additionally, Lhx2/9 morphants do not exhibit serotonin staining, indicating a lack of this neuron population (E vs. F). E and F depict only the anterior dorsal region of 96 h larvae. G. Double FISH reveals *lhx2/9* expression (magenta) in the anterior ectoderm, and co-expression with *elav* (green) in the APD, indicated by arrow heads; mesenchymal *elav* expression is also noted in the posterior embryo. H-H”. APD view of double FISH for *lhx2/9* (magenta) and *elav* (green) at 48 h. *Lhx2/9* expression frequently occurs in pairs. Asymmetric divisions generate pairs, in which one cell expresses *elav*, denoting a transition to post-mitotic neuron state (circled). I-I”. FISH (magenta) combined with EdU labeling (green) at 48 h results in co-labeling of *lhx2/9*+ cells that are actively proliferating in the anterior ectoderm (circled). DV-dorsal view. LV-lateral view.

We further investigated the relationship between *lhx2/9* and the formation of these serotonergic neurons. We show that *lhx2/9* co-localizes with *elav*+ cells in the APD at 48 h (Fig. 2G). We also see that in the APD, *lhx2/9*+ cells undergo asymmetric divisions in which one cell expresses *elav* and the other does not (Fig. 2H-H”). *Elav* expression in this territory marks post-mitotic neurons that will differentiate rather than produce additional neural precursors. Therefore, we hypothesize that the other cell in the pair, that does not express *elav*, will continue to proliferate. We also see pairs of *lhx2/9*+ cells in which neither cell expresses *elav* more posterior to the APD, which might represent cells dedicated to proliferation at that time (Fig. 2H). To test this, we next determined whether these *lhx2/9*+ cells are post-mitotic cells that are likely to be undergoing differentiation, or proliferative cells, which could serve as restricted progenitors. We again combined EdU labeling with FISH to determine whether some *lhx2/9*+ cells are dividing. We find that some *lhx2/9*+ cells are EdU positive (Fig. 2I-I”), which supports the idea that these cells are progenitors. Collectively, these data indicate that *lhx2/9* is crucial for development of serotonergic neurons because *lhx2/9*+ cells are restricted progenitors, at least some of which take on serotonergic neuron fate. This establishes a pathway of neurogenesis from multipotent progenitor to differentiated serotonergic neuron.

### Neurogenesis depends on regulatory state established spatially along the AP axis

A striking observation from this process is that the progression appears to proceed according to AP position. We observe *soxc*+ multipotent progenitors throughout the ectoderm, but only those in the anterior half of the ectoderm progress to *lhx2/9*+ restricted progenitors (Fig. S2E). Likewise, only *lhx2/9*+ cells in the APD differentiate into serotonergic neurons, and therefore become *elav*+ (Fig. 2G). We wanted to understand how neurogenesis might be tied to a GRN for AP patterning, particularly because AP patterning domains tend to be well-conserved across metazoans, although CNS morphologies are not (Lowe et al., 2003; Marlow et al., 2014; Yankura et al., 2010). We especially focused on the roles of the transcription factors *foxq2, rx*, and *six3*, in patterning distinct AP neurogenic domains. These were previously shown to be expressed in nested concentric domains along the AP axis within the ectoderm (Yankura et al., 2010). The domain of *six3* expression abuts *wnt8* expression posteriorly, while *foxq2* expression defines the most anterior ectoderm. *Rx* is expressed in the anterior half of the ectoderm, and is expressed more broadly than *foxq2*, but more anteriorly than *six3*. For simplicity, we will refer to these nested AP territories as 1-4, where 1 indicates the most anterior ectoderm and 4 the most posterior (Fig. 3A).

**Fig. 3.**
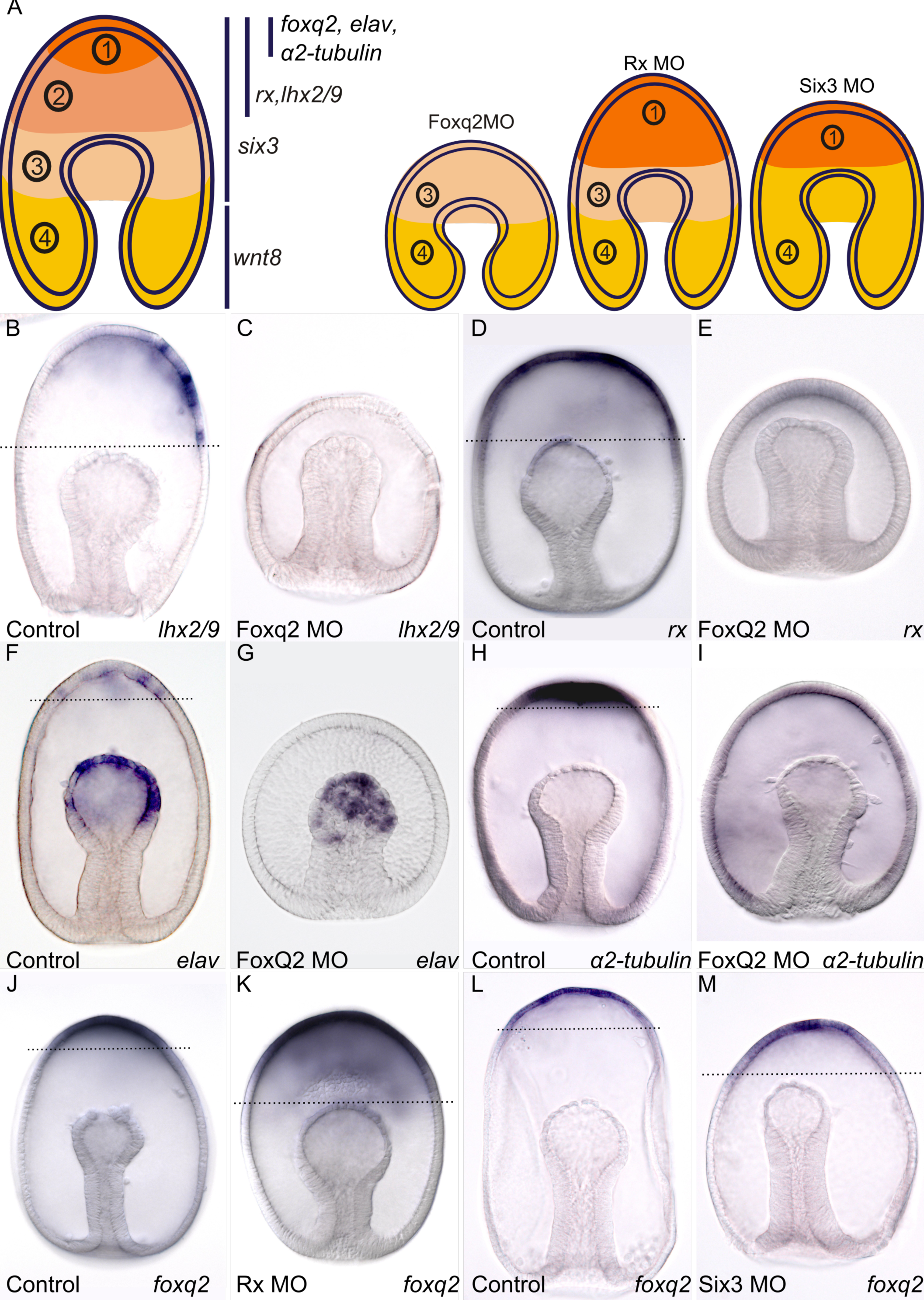
Establishment of neurogenic regions by AP domains of regulatory genes. A. Schematics depicting AP regulatory gene expression domains and alterations to these domains that occur upon knock-down of indicated transcription factors. Schematics based upon data from Figures 3 and 4, as well as our previous works (Yankura et al., 2010; Yankura et al., 2013). B-C. Normal expression of *lhx2/9* in the anterior dorsal ectoderm is lost upon perturbation of Foxq2. *Rx* expression throughout domain 2 (D) is also lost in Foxq2 morphants (E). F-I. The APD (Domain 1) expresses differentiation genes such as *elav* and *α2-tubulin*, which also require Foxq2 function. Mesodermal bulb expression of *elav* is not dependent upon Foxq2 (F-G). J. *Foxq2* expression is normally restricted to the APD. K. Rx function is required to repress *foxq2* from domain 2. Rx morphants exhibit expanded *foxq2* expression. L-M. *Foxq2* expression is not dependent upon Six3 function, but Six3 may somewhat restrict *foxq2* to domain 1.

We sought to determine the role that each of these genes has in establishing these domains. Our goal is to understand how these territories in turn may regulate neurogenesis. We first examined the role of *foxq2* in establishing domain 1. Foxq2 is required for the expression of *lhx2/9* (Fig. 3B-C) and *rx* (Fig. 3D-E) in domains 1 and 2, as well as several makers of differentiated APD specific to domain 1, including *elav* and *α2-tubulin* (Fig. 3F-I). Foxq2 therefore has a role in directing differentiation of appropriate cell type in this anterior-most ectoderm and in establishing the regulatory state of domain 2. Rx, in turn, functions to repress *foxq2* from only within domain 2, although they are co-expressed in domain 1 (Fig. 3J-K). This suggests Rx may require additional factors not present in domain 1 for repression of *foxq2*. We previously demonstrated that *six3* abuts the domain of *wnt8* expression, and one of its functions is to maintain this boundary, delineating domains 3 and 4 (Yankura et al., 2013). Wnt signaling, meanwhile, restricts the expression of *six3, lhx2/9*, and *foxq2* to their respective domains (Fig. S3C-H). Domain 1 appears unaffected in Six3 morphants, as *foxq’s* domain of expression is maintained, if not expanded (Fig. 3L-M). These morphants have, however, lost the distinct regulatory domains between *wnt8* and *foxq2* (Fig. 4A-B) as we show that *wnt8* expression now abuts or even overlaps with *foxq2* in Six3 knock-downs. Therefore, a function of both Six3 and Rx is to maintain a distinct domain 2, which is *rx*-positive, but negative for *foxq2* and *wnt8* (Fig.3A).

**Fig. 4.**
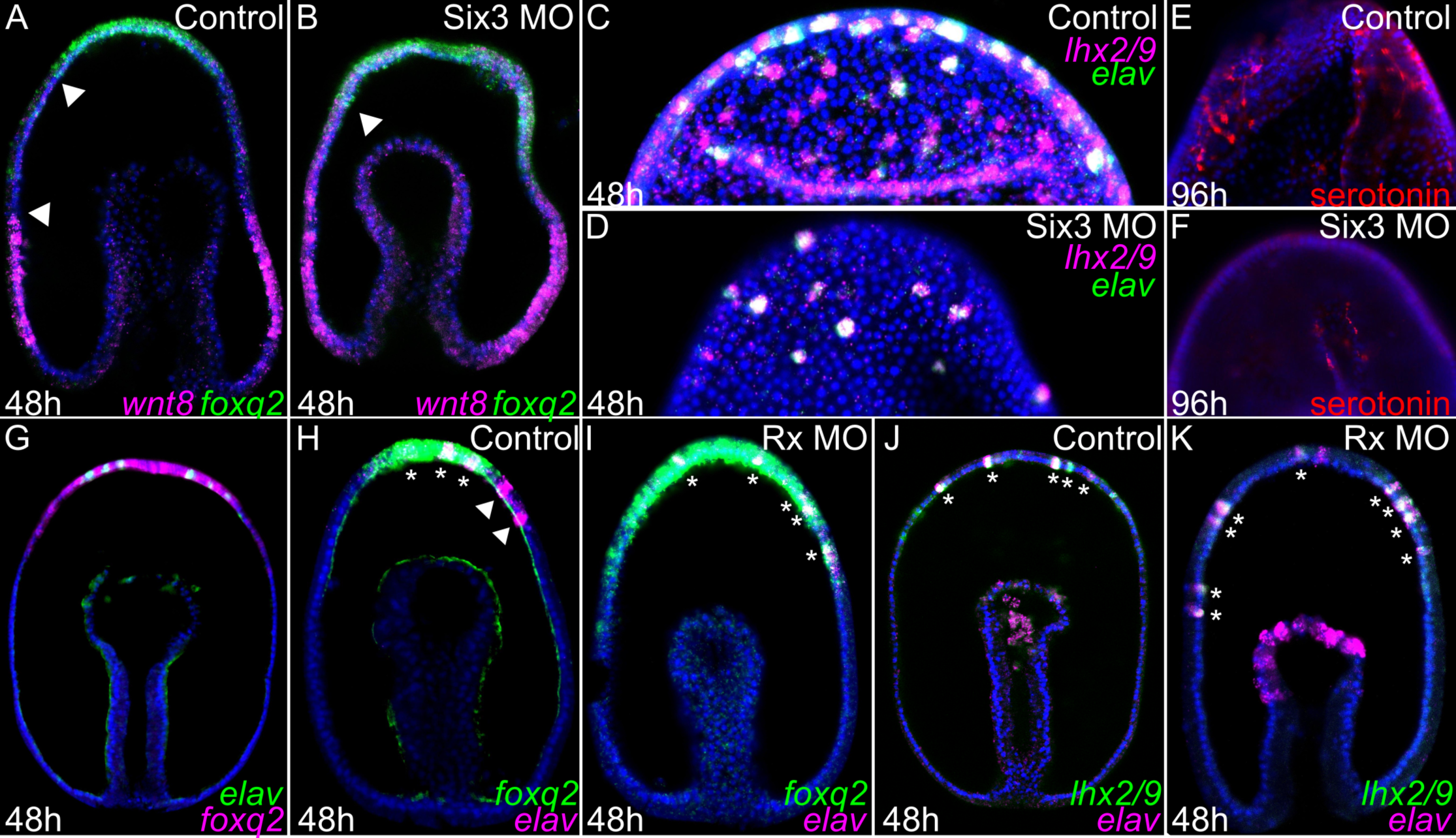
Alteration of AP domains affects the quantity and location of neurons produced. A. Two-color FISH of *wnt8* (magenta) and *foxq2* (green). *Wnt8* is restricted to domain 4, and *foxq2* to domain 1, with domains 2 and 3 in between, which express neither gene. B. Six3 morphants exhibit expansion of *wnt8* and *foxq2*, such that domain 1 and 4 abut, and unique domains 2 and 3 are lost. C. In the anterior ectoderm, *lhx2/9* (magenta) and *elav* (green) are normally co-expressed in the APD (domain 1), and pairs of cells only expressing *lhx2/9* are present more posteriorly, in domain 2. D. All *lhx2/9*+ cells also express *elav* in Six3 morphants. E-F. Anterior dorsal views of 96 h larvae. E. Control larvae have two large clusters of serotonergic neurons, indicated by immunostaining (red). F. Six3 morphants produce fewer serotonergic neurons by comparison. G. In control embryos, all ectodermal *elav* expression (green) is contained within domain 1, indicated by *foxq2* expression (magenta). H. *Lhx2/9* expression (magenta) occurs in both domains 1 and 2, therefore some *lhx2/9*+ cells are outside of the *foxq2* expression domain (green). I. In Rx morphants, *foxq2* expression is expanded and all *lhx2/9* expression is within the foxq2 domain. J. *Lhx2/9* (green) and *elav* (magenta) co-expression ordinarily occurs in domain 1, with additional *lhx2/9* only cells in domain 2. K. In Rx morphants, all *lhx2/9*+ cells co-express *elav*.

Having established, in part, the epistatic relationships that describe the initial GRN that delineates these domains, we used this knowledge to experimentally manipulate the regulatory state to determine how these territories might govern neurogenesis. We show that Six3 is not required for the correct expression of *soxc* (Fig. S3I-J). This is consistent with the broad expression of *soxc*+ cells throughout the ectoderm; i.e., they are not limited to a particular AP domain. This result also corroborates with our previous finding that the ciliary band neurons develop normally in Six3 morphants in spite of these embryos' altered morphology (Yankura et al., 2013).

We next examined the expression of *lhx2/9* in Six3 and Rx morphants. In both of these perturbations there is no longer a molecularly distinct domain 2. In Six3 morphants, in which *wnt8* expands into the anterior ectoderm, such that it now abuts the expression domain of *foxq2* (Fig. 4A-B), *lhx2/9* is still expressed, but the number of *lhx2/9*+ cells is reduced (Fig. 4C-D). In later development, the *lhx2/9*+ cells and serotonergic neurons, while still present, are also much reduced in number (Fig. 4E-F, Fig. S3K-L). This indicates that Six3 normally functions to maintain an *lhx2/9*+ progenitor population. While the more posterior *lhx2/9*+ cells normally do not express *elav* and instead can continue to proliferate in control embryos, all *lhx2/9*+ cells now co-express *elav* in Six3 morphants (Fig. 4C-D). This is presumably because all *lhx2/9*+ cells are now within domain 1, as the distinct domain 2 is lost. *Elav* is only expressed within this domain in normal embryos (Fig. 4G), suggesting that *foxq*2 could be important for serotonergic neuron cell-cycle exit and differentiation.

To test this further, we next examine the expression of *lhx2/9* in Rx morphants where *foxq2* expression expands posteriorly (Fig. 4H-I). This again places all *lhx2/9*+ cells into the *foxq2* territory, but now the domain of *foxq2* is extended. We show that, as predicted, all of these cells also express *elav* rather than maintaining a progenitor state (Fig. 4J-K). Therefore, all *lhx2/9*+ cells become *elav*+ in Six3 and Rx morphants, because they are within domain 1. This provides further evidence that *foxq2* promotes exit from proliferation, and the differentiation of restricted progenitors. As a result, the proliferating reserve of *lhx2/9*+ cells is lost in these AP domain perturbations. These results indicate, therefore, that domain 2 serves as a neural proliferation zone.

## Discussion

A critical question in developmental biology is how neural stem cells progress to their correct differentiated fate, and how this is coordinated in space and time. We show here that the sea star larva uses a simple and elegant patterning system to direct broadly localized *soxc*+proliferative cells to give rise to *lhx2/9*+ restricted progenitors. These in turn will exit their proliferative state to form serotonergic neurons in the *foxq2*+APD. Combined, our data indicates that *soxc*+ cells are multipotent progenitors in sea stars and that spatial domains along the AP axis establish regulatory states that control neural progression.

Both here, and previously (Yankura et al., 2013), we show that *soxc*+ cells are the initial precursor cells required for the eventual production of neurons in the sea star larvae. This role of Soxc in maintaining cell proliferation is most likely ancient, as its expression has recently been observed in regions of high cell division in ctenophores (Schnitzler et al., 2014). Recent work also indicates that *soxc* expressing cells contribute to the serotonergic neurons in sea urchins (Garner et al., 2015). Additionally, in the vertebrate forebrain, *sox11*, a *soxc* ortholog, facilitates the transition from multipotent progenitor to differentiating post-mitotic neuron by turning on genes such as *lhx2* (Bergsland et al., 2011). Sox11 is also needed for proper proliferation of neural progenitor cells (Wang et al., 2013). Combined, these data suggest that this early neurogenic step is likely to be conserved not only among deuterostomes, but potentially across metazoa. We had also previously correlated the presence of neurons in both the ciliary bands and the apical organ with *elav* expression (Yankura et al., 2013). Until now, we have had little knowledge of the transition state that occurs between these multipotent *soxc*+ cells and the resulting unique neural populations. Here, we demonstrate that in sea stars, *lhx2/9*+ cells are the subset of multipotent progenitor progeny that are restricted to becoming the serotonergic neurons of the apical organ, as opposed to ciliary band neurons. *Lhx2/9* promotes both proliferation of restricted progenitors and their eventual differentiation into serotonergic neurons.

An important finding from our work here is that the progression of neurogenesis is dependent on position along the AP axis. A great body of work demonstrates similarities between the AP patterning of gene expression domains within the invertebrate ectoderm and the vertebrate CNS and in particular the forebrain (Range, 2014). The purpose of such domains in the larvae of marine invertebrates has been unclear, as they do not appear use these domains to produce a complex CNS with discrete territories and cell types, as vertebrates do. Here, we explain for the first time that these expression domains define territories devoted to particular steps of neurogenesis during the development of an anterior sensory structure in sea star larvae. The entirety of this process is depicted in our model shown in Figure 5. Activating and repressive interactions between Wnt signaling, *six3, rx*, and *foxq2* result in nested gene expression of the aforementioned genes. While multipotent progenitors, marked by expression of *soxc*, are present throughout the four resulting ectodermal domains, they are only able to progress to a restricted progenitor state, indicated by expression of *lhx2/9*, in Domains 1 and 2. We demonstrate that Foxq2 provides cues needed for neural precursors to differentiate into serotonergic neurons in Domain 1. Domain 2 is a designated neural proliferation zone, as restricted progenitors are produced and continue to divide here, rather than differentiating. We find that Rx is important to the maintenance of a restricted progenitor population adjacent to the APD, as it represses *foxq2*. Additionally, we find the role of Six3 is to maintain Domain 2 by preventing both Wnt8 and Foxq2 activity from encroaching into this region. The presence and extent of these domains is crucial to ensuring that the apical organ attains a proper size by determining the number of restricted progenitors produced.

**Fig. 5.**
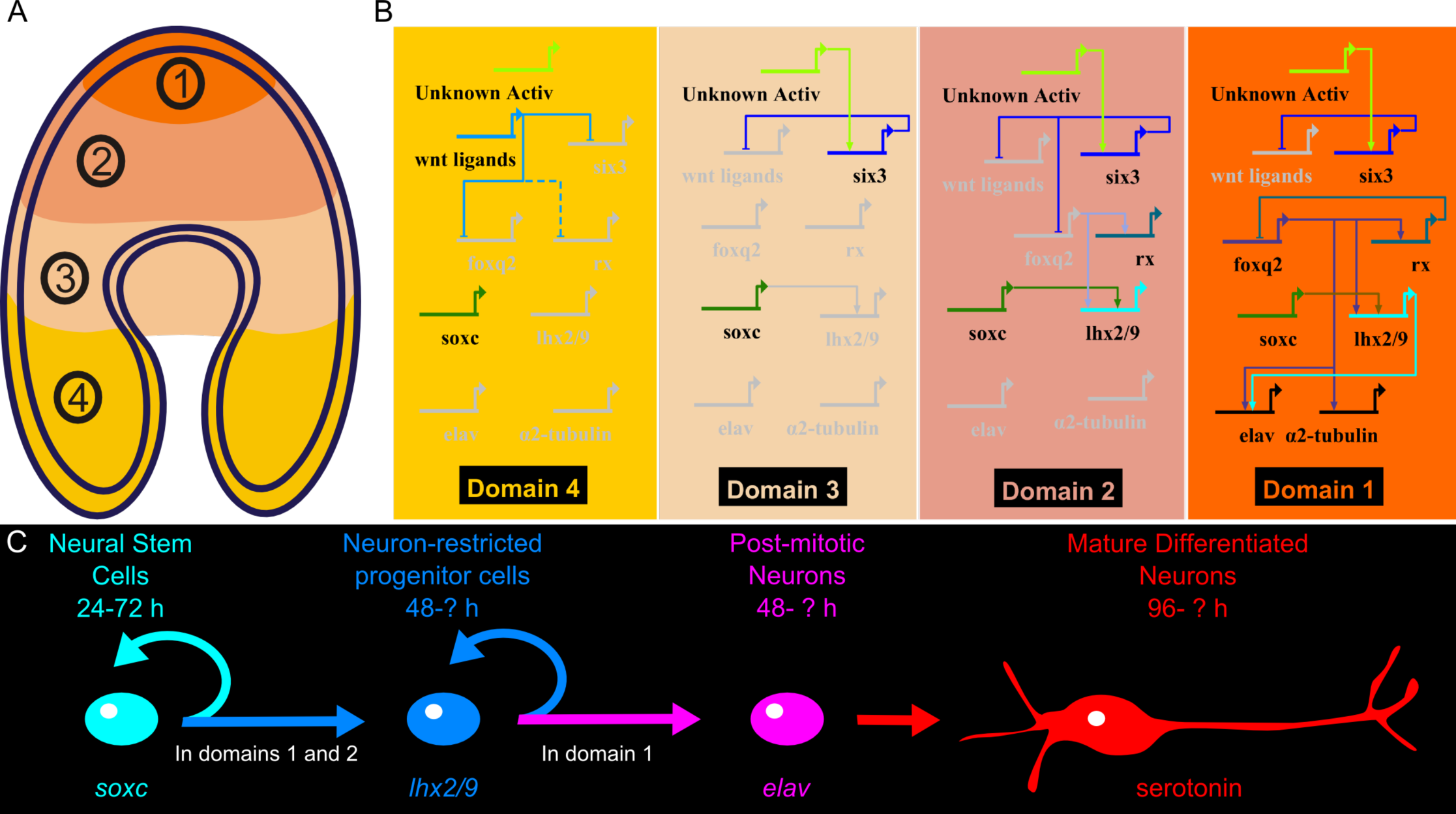
A gene regulatory network for apical organ neurogenesis. A. Schematic of AP regulatory domains in the sea star 48 h embryo. B. GRN activity is specific to each domain, resulting in different steps of neurogenesis occurring in specific regions. It is important to note that at this time, any of these regulatory inputs could be direct or indirect. CRM analysis of genes in this network have not yet been performed. C. Schematic of neurogenesis. First, restricted progenitors are produced from multipotent progenitors through asymmetric divisions. Next, post-mitotic neurons arise from asymmetric divisions of restricted progenitors. Finally, post-mitotic neurons mature into differentiated serotonergic neurons.

Interestingly, Foxq2 is able to control regulatory events in Domain 2, even though its expression is limited to Domain 1. We find that Foxq2 morphants do not express *lhx2/9* at all (Fig. 3C), rather than only missing these cells in the most-anterior region of the ectoderm. The same is true for *rx* expression (Fig. 3E). There are several possible explanations for this phenomenon. First, Foxq2 may be able to exert an effect on a Domain 2 through diffusible molecules such as the Wnt agonist, *dkk3* (Fig. S3A-B). Alternatively, this function may result from protein localization of Foxq2 in domain 2 lingering from earlier time points, as this gene is known to progressively restrict its expression domain in other echinoderms (Yaguchi et al., 2008). However, if this is the case, then progression of *lhx2/9*+ cells to serotonergic neurons must either require additional cues in Domain 1, or be prevented in Domain 2 by some unknown mechanism. Otherwise, we should observe *elav*+ cells in Domain 2. Further exploration into the regulatory states of each domain and the wiring between regulatory genes expressed in these domains will enhance our understanding of neurogenesis in the sea star and other organisms.

It had been previously suggested that Six3 must function at the top of the GRN for apical organ development in the sea urchin (Wei et al., 2009), as knock-down of this gene resulted in loss of a differentiated APD, including the loss of serotonergic neurons and apical tuft cilia. Six3 MO sea star larvae by comparison develop serotonergic neurons, although they are fewer in number. We think that this discrepancy is the result of different network wiring between *six3* and *foxq2*. While Wei and colleagues found evidence for Six3 positively regulating *foxq2* (Wei et al., 2009), we see that, in sea stars, Six3 is a likely repressor of *foxq2* (Fig. 3L and M). During mammalian telencephalon development, Six3 promotes proliferation of neural precursors by keeping these cells in an immature state and by prolonging their amplification period (Appolloni et al., 2008). As in the sea star larvae, in vertebrates, loss of Six3 does not alter the early specification of the anterior neural region; however, it does ultimately lead to truncation of the forebrain, presumably due to expansion of Wnt signaling into this region (Lagutin et al., 2003). Thus, Six3 has a highly conserved role in maintaining patterning that is tightly coupled to regulation of neural precursor proliferation.

Finally, we anticipate that evolutionary changes to the relative sizes of these domains might provide a mechanism for altering the ratio of proliferation to differentiation. There is a great diversity in neuron number among larval forms; even among echinoderms, there are potentially differences in the size of both the neural proliferative zone and the resulting apical organ. The 96 h sea star apical organ is composed of 30-50 serotonergic neurons located in two broad dorsal ganglia, while sea urchins of a comparable larval stage have only 8 serotonergic neurons, and these are restricted to a small apical pole territory (Byrne et al., 2007). We have previously compared gene expression along the AP axis of the sea star and sea urchin and suggested that while the sea urchin utilizes the same suite of regulatory genes and similar nested expression domains, the domains are extremely compacted in sea urchins compared to sea stars (Yankura et al., 2010). Study of the sea urchin reveals at least two concentric domains of gene expression in the anterior-most ectoderm; an outer ring of *six3* and an inner ring of *foxq2* (Wei et al., 2009). It is unclear whether there is a distinct ring of *rx* expression in the sea urchin ectoderm. If so, domains 2 and 3 are still much smaller in sea urchins. Meanwhile, the wide domain of *six3* expression in the sea star allows a broader region of ectoderm to be permissive to the proliferation of neuronal precursors. We find that when the functions of Six3 are blocked in the sea star, this region of proliferating restricted progenitors is lost, resulting in fewer serotonergic neurons and a restriction of these neurons to a small region in the anterior of the embryo. This phenotype is much like the normal pattern of serotonergic neurons in the sea urchin (Bisgrove and Burke, 1986; Wei et al., 2009), suggesting that changes in *six3* expression and function may have contributed to differing neural proliferative zone size and apical organ morphologies among echinoderms.

## Materials and Methods

### Embryo Culture and Microinjections

*Patiria miniata* embryos were cultured and injected as described previously (Cheatle Jarvela and Hinman, 2014). Morpholino antisense oligonucleotides were designed by GeneTools. Sequences are available in SI. Images depicting morpholino injected embryos are representative of phenotypes observed in at least three experiments, each containing 30-50 morphant embryos. Control embryos are siblings of corresponding gene knock-down embryos and were injected with 400μM standard control morpholino construct (GeneTools).

### BAC Construct Development and Injections

The *Patiria miniata* BAC library was screened with a *soxc*-specific RNA probe. Five possible *soxc* containing BACs were selected and sequenced. Two were found to have a 150Kbp insert with approximately 75Kbp of sequence up and downstream of the *soxc* transcript start and standard protocols were used to recombineer a GFP sequence into the *soxc* single exon in each. Sequencing was used to confirm successful integration (Yu et al., 2000). Constructs were prepared and injected as described previously (Hinman et al., 2007).

### Whole Mount In Situ Hybridization (WMISH) and Fluorescent In Situ Hybridization (FISH)

WMISH was performed as previously described (Hinman et al., 2003; Yankura et al., 2010) using digoxigenin-or dinitrophenol labeled antisense RNA probes. FISH embryos were mounted in Slowfade media (Life Technologies) and imaged by confocal microscopy with a Carl Zeiss LSM-510 Meta DuoScan Inverted Confocal Microscope.

### Immunofluorescence

Embryos were fixed as described for WMISH and stained with rabbit anti-serotonin (Sigma) and in some cases also anti-GFP (Pierce). Further detail is provided in the SI.

### EdU Labeling

EdU labeling was performed using the Click-It Plus EdU 488 and 647 Imaging Kits (Life Technologies), with some modifications to the manufacturer's instructions, as described in the SI. All EdU labeled embryos were treated with EdU reagent for 15 minutes. For pulse-chase experiment, EdU was subsequently flushed out with artificial sea water, and embryos were incubated an additional 30 minutes.

## Acknowledgements

We thank Dr. Greg Cary and Minyan Zheng for critical reading of this manuscript and fruitful discussions; Ping Dong and Eric Davidson for assistance in construction of the Soxc-GFP BAC; David McClay for kindly providing the dominant-negative cadherin construct; and Marinus Inc. and Pete Halmay and Pat Leahy for animal collection.

## Author Contributions

ACJ, KAY, and VFH devised the study, performed the experiments, and analyzed the results. ACJ and VFH wrote the manuscript.

## Funding

This work was partially supported by National Science Foundation Grant 0844948 (to V.F.H.).

## References

Appolloni, I., Calzolari, F., Corte, G., Perris, R. and Malatesta, P. (2008). Six3 Controls the Neural Progenitor Status in the Murine CNS. Cereb. Cortex 18, 553–562.

Bergsland, M., Ramsköld, D., Zaouter, C., Klum, S., Sandberg, R. and Muhr, J. (2011). Sequentially acting Sox transcription factors in neural lineage development. Genes Dev. 25, 2453–2464.

Bisgrove, B. W. and Burke, R. D. (1986). Development of Serotonergic Neurons in Embryos of the Sea Urchin, Strongylocentrotus purpuratus. Dev. Growth Differ. 28, 569–574.

Byrne, M., Nakajima, Y., Chee, F. C. and Burke, R. D. (2007). Apical organs in echinoderm larvae: insights into larval evolution in the Ambulacraria. Evol. Dev. 9, 432–445.

Cheatle Jarvela, A. M. and Hinman, V. (2014). A Method for Microinjection of Patiria minata Zygotes. J. Vis. Exp. JoVE.

Chee, F. and Byrne, M. (1999). Development of the Larval Serotonergic Nervous System in the Sea Star Patiriella regularis as Revealed by Confocal Imaging. Biol. Bull. 197, 123–131.

Garner, S., Zysk, I., Byrne, G., Kramer, M., Moller, D., Taylor, V. and Burke, R. D. (2015). Neurogenesis in sea urchin embryos and the diversity of deuterostome neurogenic mechanisms. Dev. Camb. Engl.

Guillemot, F. (2007). Spatial and temporal specification of neural fates by transcription factor codes. Development 134, 3771–3780.

Hinman, V. F., Nguyen, A. T. and Davidson, E. H. (2003). Expression and function of a starfish Otx ortholog, AmOtx: a conserved role for Otx proteins in endoderm development that predates divergence of the eleutherozoa. Mech. Dev. 120, 1165–1176.

Hinman, V. F., Nguyen, A. and Davidson, E. H. (2007). Caught in the evolutionary act: precise cis-regulatory basis of difference in the organization of gene networks of sea stars and sea urchins. Dev. Biol. 312, 584–595.

Holland, L. Z., Carvalho, J. E., Escriva, H., Laudet, V., Schubert, M., Shimeld, S. M. and Yu, J.-K. (2013). Evolution of bilaterian central nervous systems: a single origin? EvoDevo 4, 27.

Kempf, S. C., Page, L. R. and Pires, A. (1997). Development of serotonin-like immunoreactivity in the embryos and larvae of nudibranch mollusks with emphasis on the structure and possible function of the apical sensory organ. J. Comp. Neurol. 386, 507–528.

Kohwi, M. and Doe, C. Q. (2013). Temporal fate specification and neural progenitor competence during development. Nat. Rev. Neurosci. 14, 823–838.

Lagutin, O. V., Zhu, C. C., Kobayashi, D., Topczewski, J., Shimamura, K., Puelles, L., Russell, H. R. C., McKinnon, P. J., Solnica-Krezel, L. and Oliver, G. (2003). Six3 repression of Wnt signaling in the anterior neuroectoderm is essential for vertebrate forebrain development. Genes Dev. 17, 368–379.

Lowe, C. J., Wu, M., Salic, A., Evans, L., Lander, E., Stange-Thomann, N., Gruber, C. E., Gerhart, J. and Kirschner, M. (2003). Anteroposterior Patterning in Hemichordates and the Origins of the Chordate Nervous System. Cell 113, 853–865.

Marlow, H., Tosches, M. A., Tomer, R., Steinmetz, P. R., Lauri, A., Larsson, T. and Arendt, D. (2014). Larval body patterning and apical organs are conserved in animal evolution. BMC Biol. 12, 7.

Miyagi, S., Kato, H. and Okuda, A. (2009). Role of SoxB1 transcription factors in development. Cell. Mol. Life Sci. CMLS 66, 3675–3684.

Nakajima, Y., Humphreys, T., Kaneko, H. and Tagawa, K. (2004a). Development and neural organization of the tornaria larva of the Hawaiian hemichordate, Ptychodera flava. Zoolog. Sci. 21, 69–78.

Nakajima, Y., Kaneko, H., Murray, G. and Burke, R. D. (2004b). Divergent patterns of neural development in larval echinoids and asteroids. Evol. Dev. 6, 95–104.

Price, D., Jarman, A., Mason, J. and Kind, P. (2011). Neurogenesis: Generating Neural Cells. In Building Brains, pp. 91–117. John Wiley & Sons, Ltd.

Range, R. (2014). Specification and positioning of the anterior neuroectoderm in deuterostome embryos. Genes. N. Y. N 2000 52, 222–234.

Schnitzler, C. E., Simmons, D. K., Pang, K., Martindale, M. Q. and Baxevanis, A. D. (2014). Expression of multiple Sox genes through embryonic development in the ctenophore Mnemiopsis leidyi is spatially restricted to zones of cell proliferation. EvoDevo 5, 15.

Sinigaglia, C., Busengdal, H., Leclère, L., Technau, U. and Rentzsch, F. (2013). The bilaterian head patterning gene six3/6 controls aboral domain development in a cnidarian. PLoS Biol. 11, e1001488.

Wang, Y., Lin, L., Lai, H., Parada, L. F. and Lei, L. (2013). Transcription factor Sox11 is essential for both embryonic and adult neurogenesis. Dev. Dyn. Off. Publ. Am. Assoc. Anat. 242, 638–653.

Wei, Z., Yaguchi, J., Yaguchi, S., Angerer, R. C. and Angerer, L. M. (2009). The sea urchin animal pole domain is a Six3-dependent neurogenic patterning center. Dev. Camb. Engl. 136, 1179–1189.

Yaguchi, S., Yaguchi, J., Angerer, R. C. and Angerer, L. M. (2008). A Wnt-FoxQ2-nodal pathway links primary and secondary axis specification in sea urchin embryos. Dev. Cell 14, 97–107.

Yankura, K. A., Martik, M. L., Jennings, C. K. and Hinman, V. F. (2010). Uncoupling of complex regulatory patterning during evolution of larval development in echinoderms. BMC Biol. 8, 143.

Yankura, K. A., Koechlein, C. S., Cryan, A. F., Cheatle, A. and Hinman, V. F. (2013). Gene regulatory network for neurogenesis in a sea star embryo connects broad neural specification and localized patterning. Proc. Natl. Acad. Sci. 110, 8591–8596.

Yu, D., Ellis, H. M., Lee, E.-C., Jenkins, N. A., Copeland, N. G. and Court, D. L. (2000). An efficient recombination system for chromosome engineering in Escherichia coli. Proc. Natl. Acad. Sci. 97, 5978–5983.

